# Common practice solvent extraction does not reflect actual emission of a sex pheromone during butterfly courtship

**DOI:** 10.1101/270462

**Authors:** Bertanne Visser, Ian A. N. Dublon, Stéphanie Heuskin, Florent Laval, Paul M. B. Bacquet, Georges Lognay, Caroline M. Nieberding

## Abstract

Olfactory communication can be of critical importance for mate choice decisions. Lepidoptera are key model systems for understanding olfactory communication, particularly considering sex pheromone signaling in the context of sexual selection. Solvent extraction or rinsing of pheromone-producing structures is a widespread method for quantifying sex pheromones, but such measures reflect what is stored and may not represent what is actually emitted by an individual during courtship. Here, we address this point for the first time by quantifying the components of the male sex pheromone (MSP) of interacting *Bicyclus anynana* butterflies, a species for which much information is available onthe role played by MSPs in affecting mating success. Using headspace sampling during courtship and solvent extraction after completion of experiments using the same males, we were able to track individual traits. Our results show that solvent extracts do not reflect quantities of MSP components emitted by live butterflies. We further show that MSP amounts obtained using headspace sampling correlated with male mating success, but solvent extracts did not. Our results further strongly suggest that males actively control MSP emission when faced with increased male-male competition. Common practice solvent extracts may thus not serve as an adequate proxy for male sex pheromone signaling as they are perceived by choosy females. Our study serves as a proof of principle that quantification of male sex pheromone components depends on the method of collection, which could apply to many other insects using short-range chemical signals. This affects our understanding of how sexual selection shapes the evolution of sexually-selected chemical traits.

## Introduction

Sexual selection was first defined by Charles Darwin and Alfred Russel Wallace as a type of natural selection where access to reproduction depends on a specific part of the environment, i.e. the other sex (Darwin, 1859; Wallace 1892). Sexual selection can be a major driver shaping the evolution of secondary sexual traits and can also lead to the evolution of reproductive isolation (Boughman, 2001; Panhuis et al., 2001). Whilst sexual selection research has had a large focus on morphological, visual, and acoustic traits, many organisms interact mostly through chemical signals called sex pheromones (Smadja & Butlin, 2009; Wyatt, 2014). Sex pheromones can be critical for reproductive success, because these signals can convey information on the location, quality, sex, and species identity of potential mates (Johansson and Jones, 2007; Karlson and Lüscher, 1959; Wyatt, 2014) and can be involved in reproductive isolation (Bacquet et al, 2015; Wyatt, 2014). How sexual selection shapes the evolution of sex pheromones remains largely unclear, however, and different schools of thought assume that sexual selection produces either stabilizing selection on the presence and amount of chemical compounds, or directional evolution for increased amounts of specific compounds that are preferred by the other sex (Groot et al., 2016).

Lepidoptera have become important model organisms in studies on sexual selection of olfactory communication (Wyatt, 2014). After identification of the first sex pheromone in the silk moth *Bombyx mori* (Karlson and Lüscher, 1959), early work on sexually selected olfactory signals focused on female moths that release remarkably long-range pheromone plumes to attract conspecifics (Greenfield, 1981). Female moth mating signals tend to be similar among closely related species and sexual communication plays a major role in species recognition (Groot et al 2016). However, many male moths and butterflies also produce sex pheromones, typically during courtship, and emitted at close range (Andersson, Borg-Karlson, Vongvanich, & Wiklund, 2007; Birch, Poppy, & Baker, 1990; Nieberding et al., 2008; Phelan & Baker, 1987; Sappington & Taylor, 1990). It is these close-range signals that are expected to play a major role in mate-choice decisions, yet chemical communication by male moths and butterflies has received relatively little attention (Heuskin et al., 2014; Nieberding et al., 2008).

Quantification of chemical signals has relied heavily on solvent extraction, a method by which pheromone-producing structures are removed and subsequently soaked or rinsed in a solvent (Darragh et al., 2017; Wyatt, 2014). A recent overview of pheromone signaling in Lepidoptera reported that at least 85% of studies used solvent extraction/rinsing of pheromone-producing glands to quantify pheromone levels (Umbers et al., 2015). Results of this common practice technique have been used as a proxy for both pheromone synthesis and release (Foster et al., 2018), but what is contained within a pheromone-producing structure may not reflect what is actually emitted during courtship. Indeed, Byrne et al. (1975) already recognized that solvent extraction may not reflect pheromone emission and designed a headspace sampling method for insects using absorption on Porapak Q. This method was then successfully used to measure sex pheromones of several lepidopteran species (Kuwahara 1979; Hirai 1980). What is striking is that most studies using Byrne et al’s method found that pheromone quantities or ratios were not similar when determined by headspace sampling or solvent extracts (Byrne et al 1975; Roelofs et al 1975; Hill et al 1975; but see Toth and Buser, 1992). The need to collect pheromones from air rather than tissue extracts was thus already clear in the 70s (Hill et al 1975; Sanders and Weatherston 1976; Percy et al 1971; Cross et al 1976). Soaking or rinsing sex pheromone-producing tissues may thus not reliably quantify olfactory sexual signals as they are emitted and perceived by the choosy sex during courtship behavior.

To understand how short-range pheromone signaling affects mate-choice decisions and sexual selection, it is essential to accurately quantify signals as they are perceived by the other, choosy sex, because it is this information that sexual selection acts upon. We hypothesized that common practice solvent extraction could blur sex pheromone quantification and as such limit our understanding of how sexual selection affects chemical sexually-selected traits. We used the butterfly *Bicyclus anynana* (Lepidoptera: Nymphalidae) as a model system, because this species is one of the best studied butterflies with regard to sexual, including olfactory, communication (Bacquet et al., 2015; Brakefield et al., 2009; Costanzo and Monteiro, 2007; Dion et al., 2016; Heuskin et al., 2014; Nieberding et al., 2008, 2012, 2018; Nieberding and Holveck, 2017; Prudic et al., 2011; San Martin et al., 2011; van Bergen et al., 2013; Nieberding et al., 2018; Westerman et al., 2012, 2014). In *B. anynana*, males compete for access to females and perform a stereotyped courtship sequence during which the sex pheromone is emitted by wing pheromone-producing structures called androconia (Figure 1). The male sex pheromone (“MSP” hereafter) is composed of three active components: (Z)-9-tetradecenol (Z9-14:OH; MSP1), hexadecanal (16:Ald; MSP2), and 6,10,14-trimethylpentadecan-2-ol (MSP3). Females do not emit these MSP components, but have the olfactory receptors to perceive these three MSP components on their antennae (Heuskin et al., 2014; Nieberding et al., 2008). Male wings produce larger amounts of MSP3 than MSP1 or MSP2 (Nieberding et al., 2012). Males with artificially reduced MSP production suffer from reduced mating success (Costanzo and Monteiro, 2007; Nieberding et al., 2008) and MSP composition was found to be a reliable indicator of male identity, level of inbreeding, as well as age (Nieberding et al., 2012; van Bergen et al., 2013). Females were thus shown to use variation in absolute and relative amounts of these three components in deciding with whom to mate.

**Figure 1:**
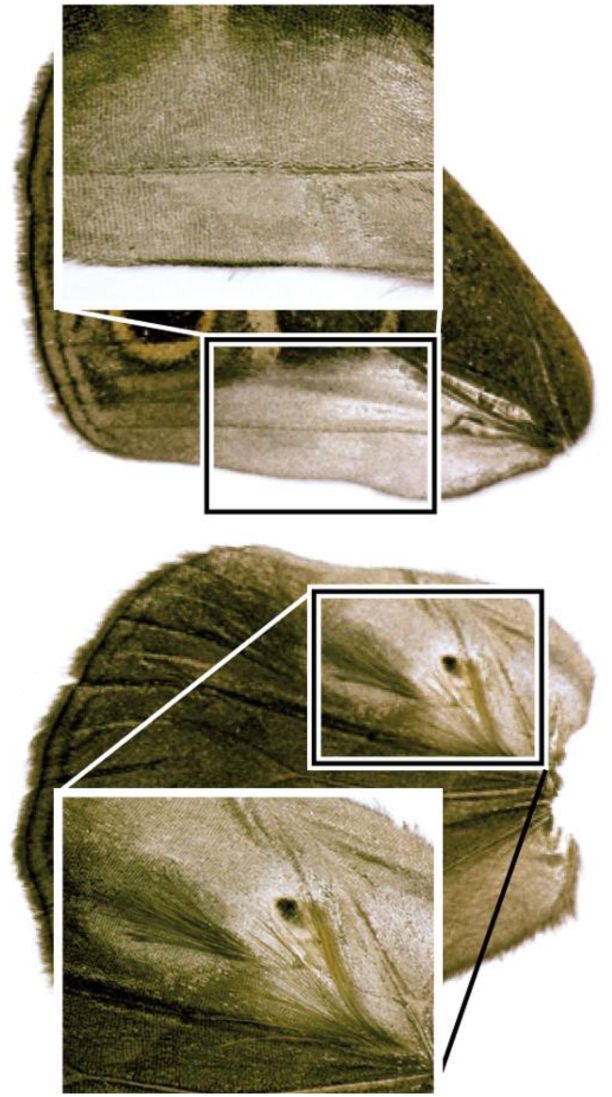
Male ventral forewing containing sex scales (top) and dorsal hindwing containing both sex scales and androconia (bottom)

To test whether solvent extracts provide an accurate estimate of chemical signals emitted during male butterfly courtship, we used an entrainment chamber to compare MSP amounts emitted in the air by live, courting males, i.e. headspace extracts, to solvent extracts of the same individuals after completion of experiments. We further determined whether amounts obtained through headspace and solvent extraction were correlated with male mating success and if males actively control the emission of MSP. We discuss how methodological choices affect our understanding of sexual selection on olfactory communication in a butterfly species.

## Materials and methods

### *Model organism* Bicyclus anynana

An outbred laboratory population of *B. anynana* was established at the Université catholique de Louvain (Belgium) in 2012 from an existing laboratory population that was established in 1988 from over 80 gravid field-caught females in Malawi, Africa. Larvae were reared on maize (*Zea mays mays*) and adults fed bananas (*Musa acuminata*) *ad libitum*. Population sizes were maintained at ~400 to 600 adults for each generation to preserve high levels of heterozygosity (Brakefield et al., 2009). Experiments were performed on individuals reared in a climate chamber under a standardized temperature regime at 26.0 ±2.0°C, a relative humidity of 70 ±15% and a photoperiod of 12:12 L:D, representing the tropical wet season under natural conditions. Sexes were separated on the day of emergence and virgin males and females between 7 to 10 days and 4 to 6 days of age, respectively, were used for experiments, because at these ages males produce significant amounts of all 3 active MSP components and females are sexually mature and show high mate preference and selectivity (Nieberding et al, 2012).

### Experimental set-up for headspace sampling

A custom-built headspace entrainment arena with a capacity of 1.8 L (Pierre E. ltd., Vilvoorde, Belgium) was used to collect volatile chemical components produced by live *B. anynana* males. Custom-made sorbent cartridges were prepared with 60 mg Tenax-TA 20/35 sorbent (04914, Grace Davidson Discovery Science, IL) in glass tubes (Figure 2). Sorbent cartridges were coupled to Teflon tubings (BOLA PTFE 8 mm i.d.) at both sides with one side facing the arena and the other side facing an air pump (Escort ELF Personal Air Sampling Pump, Zefon International Inc., Florida USA) operating at 0.8 L min^-1^. Airflow was verified prior to connection to the system using a digital flowmeter (MesaLabs Bios Defender 520, Colorado, USA). Sorbent cartridges were further cleaned by flushing with 1.5 ml of 90:10 v/v mixture of n-hexane and diethyl ether and left to dry before each experiment. Prior to use the whole entrainment system was thoroughly cleaned.

**Figure 2:**
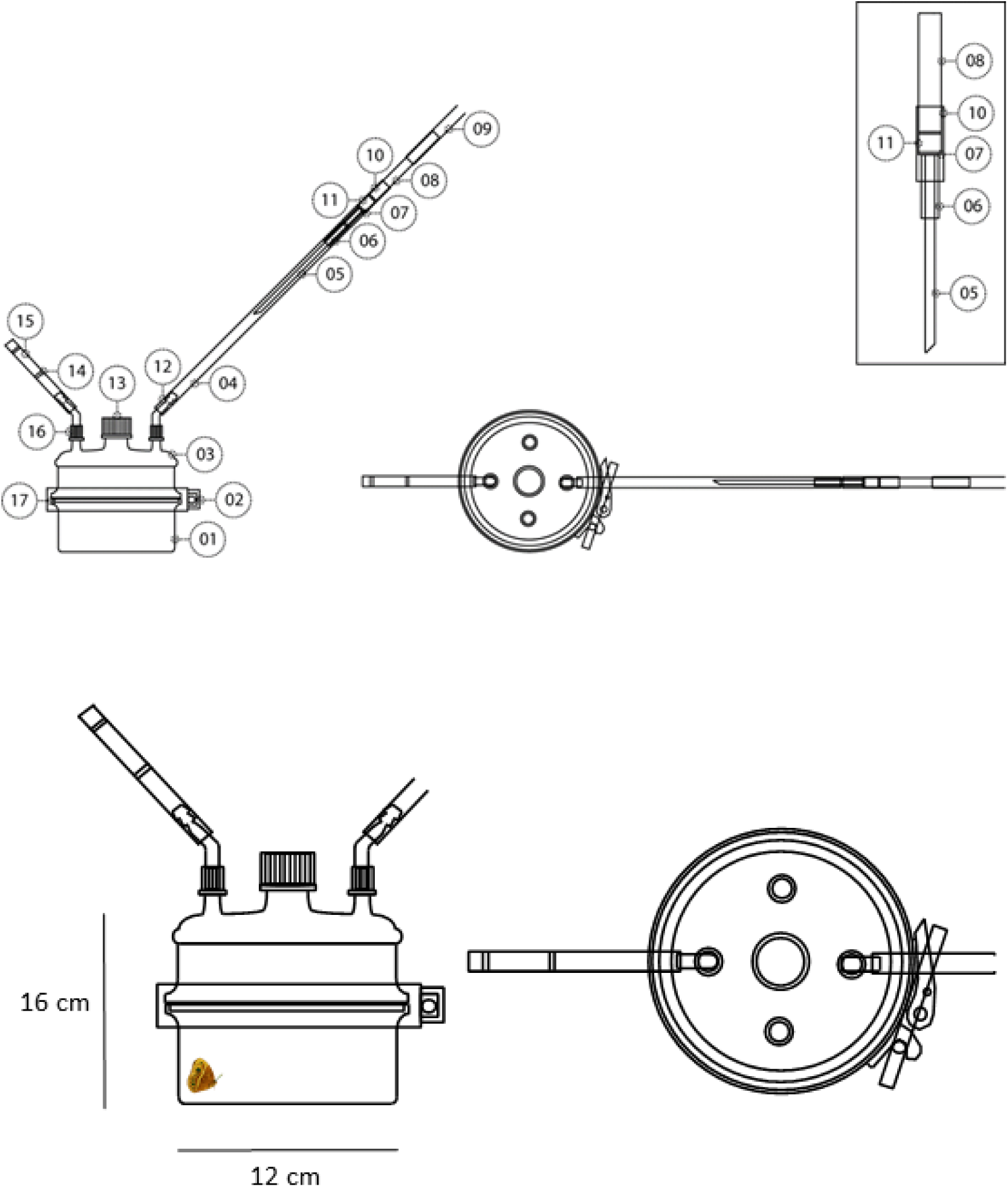
Top: Headspace entrainment area. Left: side view (with two smaller ports omitted from the lid). Right: viewed from above showing all ports. Labels: 01: culture flask injection head; 02: stainless steel band; 03: corresponding culture flask lid with added ports; 04: 8mm PTFE tubing; 05: 3mm glass with 45 degree cut; 06: 5.5 mm outer diameter PTFE; 07: 7mm outer diameter PTFE; 08: 7mm outer diameter glass; 08: 8mm PTFE tubing; 10: metal grid; 11: Tenax-TA 20/35; 12: silicone rubber sealed plastic hose connector; 13 GL45 centralized screwthread with cap; 14: glass wool plug; 15: activated dry carbon; 16: GL14 screwthread and cap with aperture; 17: PTFE ‘O’ ring. Inset: magnified annotated sorbent cartridge design. Bottom: magnified view from side and above.

### Behavioral observations

To determine mating success during experiments, male abdominal tips were dusted with different colors of a U.V. fluorescent powder dye (‘rodent-tracking’ fluorescent dust, chartreuse “TP35” Radiant Color NV, Houthalen, Belgium) to allow tracking of copulation events through dust transfer between genitalia (i.e. female abdomens will contain fluorescent dye if mating occurred)(Joron and Brakefield, 2003). We produced three treatments with increasing male-biased sex ratio: 1:3, 1:1 and 3:1 males to females, using different virgin males and females, with 14, 15 and 14 replicates, respectively. The entrainment chamber was headspace sampled over a 22.5 hour period. To link pheromone emission to male activity, behaviors were observed and recorded using the program The Observer v. 5.0 (Noldus, Hilversum, the Netherlands). Recorded behaviors included the number and duration of general activities (walking, flying), as well as the number and duration of courtship behaviors (i.e. courtship sequences that included typical male wing fluttering behaviors; Nieberding et al 2008). Behavioral observations started around 14:00. Male activity was examined during 15 minutes at the start of the experiment, and for another 15 minutes one hour later (starting at 15:00). Courtship activity takes place during the entire daylight phase, but courtship activity peaks in the afternoon and observations of 30 minutes during peak activity provide representative quantification (Nieberding et al. unpublished data). Moreover, previous work suggests that repeatability of courtship activity is improved when daily observations take place at the same time of day, as opposed to repeated measures during the day, because variation in activity varies about 10-fold throughout a day (Nieberding et al, unpublished data). To avoid stress during the entrainment period, a non-sterile cotton wool segment (~60 mm x ~40 mm) containing ~5 ml of cane sugar solution diluted in water (5 g in 200ml^-1^) was added to the entrainment arena. This allowed *ad libitum* feeding without volatile contamination. Control entrainments in which no insects were added to the arena were also performed to verify the absence of MSPs (levels < LOD). After the entrainment was terminated, female genital regions were viewed under UV light at 365nm (18W Blacklight-Blue F18W/T8/BLB, Havells-Sylvania, Antwerp Belgium) to determine if mating had occurred during the 22.5 hour entrainment period. Males were then collected and frozen at −80 °C, after which wings were removed and used for MSP quantification by solvent extraction (see below).

### MSP quantification using headspace and solvent extracts

After each entrainment experiment, sorbent cartridges were eluted twice with 200 μl 90:10 v/v n-hexane-diethyl ether. The more polar solvent ether was added to recover all trapped products leading to total desorption. Ten μl of trans-4-tridecenyl acetate was then added to the elution solvent to provide an internal standard with a final concentration of 5ng μl^-1^. This enabled direct comparison with obtained peak areas. As elution from the cartridge was expected to be less than the full solvent volume applied, 10 μl of a second standard (C10 butylbenzene; final concentration of 1 ng μl^-1^) was added directly prior to cartridge elution. Analysis of the butylbenzene peak area within a complete 220 μl solvent volume enabled us to quantify actual solvent loss in every elution. GC analyses were carried out on an Agilent GC7890A gas chromatograph fitted with a flame ionisation detector (Agilent Technologies, Belgium) and a splitless injector at 250 °C. A 30m × 0.32 mm DB-5 (df=0.2 μm) column (Agilent, 19091J-413) was used with H_2_ as the carrier gas at a constant flow of 30 ml.min^-1^. The temperature program was as follows: initial temperature of 75°C for 3 minutes which was then programmed to 220°C at 20°C min^-1^ until 300 °C at 30°C min^-1^ with a final hold of 7 min. The FID was maintained at 250°C. Injections were made using a 7693 ALS autosampler (Agilent), injecting 1μl. All acquisitions and integrations were examined with GC Chemstation B.04.03-SP2 (Agilent). Solvent extractions were performed according to Nieberding et al (2008) and Heuskin et al (2014). Briefly, MSP components were extracted by placing one fore- and one hind-wing of each male in 350ul n-hexane, which contained an internal standard (trans-4-tridecenyl acetate at 10 ng ul^-1^), for 10 min. Separations were carried out in the aforementioned chromatographic conditions. This allowed for a direct comparison between ‘on-wing’ MSP levels and headspace MSP collections.

### Statistics

Statistical analyses were done using R 3.3.1 (R Development Core Team, 2016) via the RStudio Desktop v0.99.903 (RStudio Inc., Boston, Massachusetts, USA). We used a linear model to test for a correlation between MSP amounts obtained using headspace sampling and solvent extracts (for MSP1 and MSP3 separately). MSP1 and MSP3 quantities were further compared between methods using t-tests. All replicates were used for these tests (n = 43). A linear mixed effects model (GLMM; lme4 package) was then used to test for the effect of mating number and sex ratio on MSP quantities with the following structure: *Y ~ sex ratio* (fixed) + *mating number* (fixed) + *sex ratio x mating number* + *male age* (random) + *female age* (random). All replicates were used for this model (n = 43; MSP amounts and behavioral activity were averaged by male numbers when 3 males were present in a single replicate). To test for the effect of courtship behaviors, general movements and sex ratio on MSP quantities the following linear mixed effects model was fitted: *Y ~ sex ratio* (fixed) + *log courtship behaviors* (fixed) + *log general movements* (fixed) + *log courtship behaviors x log general movements* (fixed) + *male age* (random) + *female age* (random). For the latter model, only experiments where a single male was present were used (n = 28) in order to have exact data per individual for both behaviors and MSP components. Full models went through model simplification to obtain minimal adequate models (i.e. non-significant terms were sequentially removed).

## Results

### Do tissue extracts reflect olfactory signals emitted during courtship?

We aimed to determine if one of the most commonly used methods in studies on olfactory communication, tissue extraction/rinsing in a solvent, reflects olfactory signals as they are emitted in the air. To test this, quantities of MSP components emitted in the air during courtship were determined using headspace sampling and compared with solvent extracts in hexane of the same individual after experiments ended. Average MSP amounts per male differed strongly between methods for MSP2 and MSP3, but not for MSP1. For MSP1, MSP quantities found on the wing at the end of behavioral experiments were similar to what was emitted by males during a day of courtship activity (solvent extract mean ± 1SE: 3.3 ± 0.06 μg; headspace extract mean ± 1SE: 2.9 ± 0.3 μg; t-test, t = −0.73; df = 83.39; p = 0.47). In contrast, MSP3 was detected in the air at a concentration almost ten times lower than what was extracted from the wings (solvent extract mean ± 1SE: 11.7 ± 0.8 μg; headspace extract mean ± 1SE: 1.2 ± 0.2 μg; t-test t = −8.42; df = 43.21; p < 0.0001). Contrary to what solvent extracts tell us, MSP3 is thus not the most abundant chemical emitted by courting *B. anynana* males. We further found a correlation in MSP1 and MSP3 quantities between solvent extracts and headspace sampling of individual males (MSP1: [inline], F_1,41_ = 12.95, p < 0.001; MSP3: [inline], F_1,41_= 9.25, p < 0.01; Figure 3). Surprisingly, we did not find any MSP2 in headspace samples, while MSP2 was present in typical amounts in solvent extracts (solvent extract mean ± 1SE: 1.2 ± 0.06 μg)(Heuskin et al., 2014; Nieberding et al., 2008, 2012).

**Figure 3:**
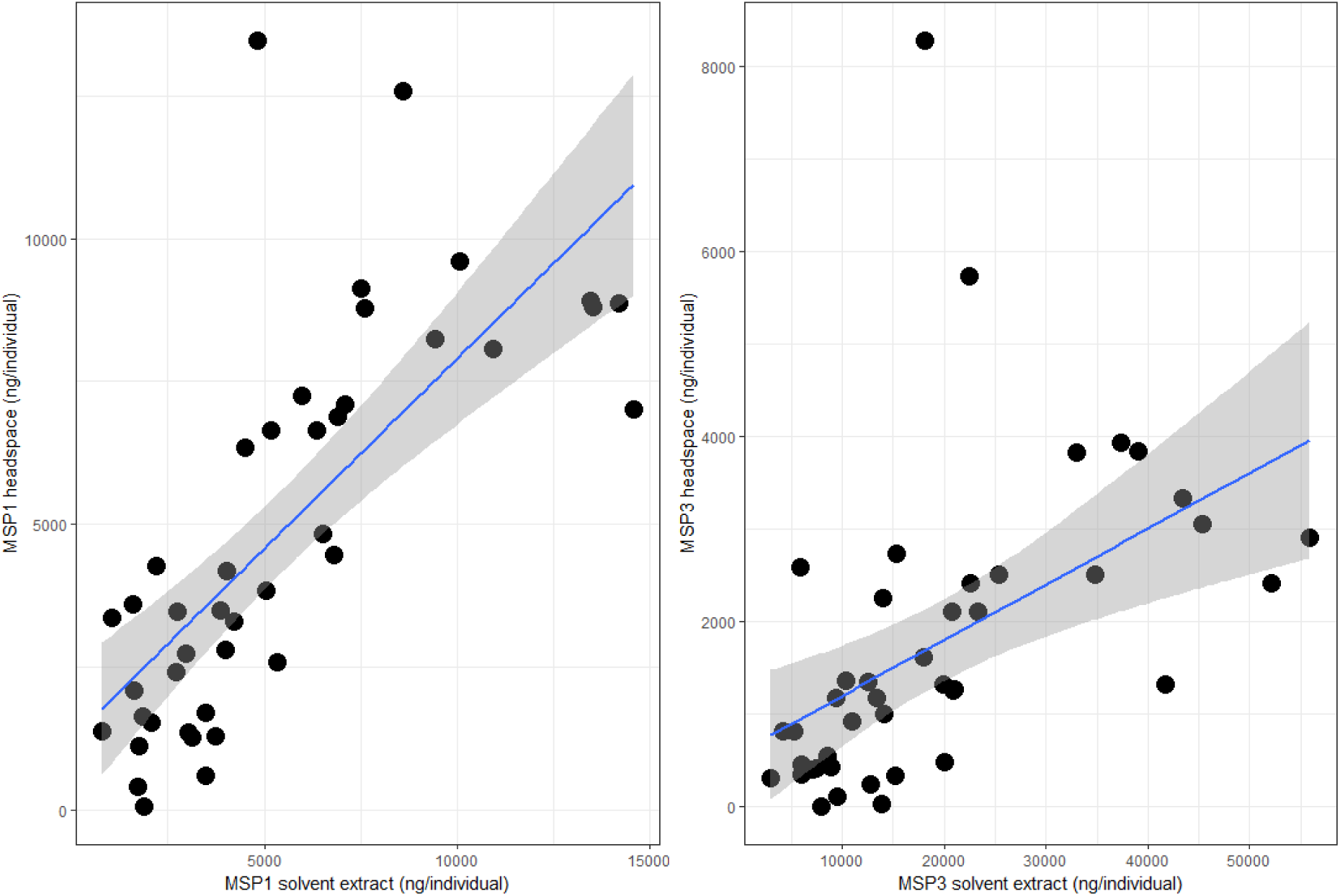
Correlation between MSP1 (left) and MSP3 (right) amounts (in ng/individual) and confidence intervals obtained by headspace sampling live butterflies during 22.5 hours (Y axis) or wing tissue solvent extraction after completion of behavioral experiments (X axis). N = 43.

### Do MSP amounts contribute to mating success?

We produced different sex ratios to manipulate the level of male-male competition: female-biased, equal, or male-biased sex ratios (1:3, 1:1 and 3:1, male:female). The relative number of mated males (i.e. the number of matings divided by the number of males within one treatment) decreased with increasing male-biased sex ratios, from 1.4 (+/- 0.17, 1SE) relative matings under female-bias to 0.6 (+/- 0.05, 1SE) relative matings under male-bias (R_adj_ = 0.31; F_1,41_= 19.9; p < 0.001). As expected, increased male-male competition was thus associated with increasing male-biased sex ratio (Holveck et al., 2015; Nieberding and Holveck, 2017; Nieberding and Holveck, 2018). We further expected that increasing male-male competition would induce males to produce and/or emit more MSP, as MSP is a trait under directional sexual selection (Nieberding et al 2012, Van Bergen et al 2013). We found that amounts of MSP1 and MSP3 components quantified in the air using headspace sampling increased with the number of matings (Tables 1 and 2; Supplementary Figure 1). In contrast, MSP1 and MSP3 components quantified by solvent extraction did not covary with number of matings (Tables 1 and 2; Supplementary Figure 1). We could not compare the role of MSP2 as the latter was not detected in headspace extracts. Hence, mating success is associated to larger amounts of MSP components as quantified in headspace extracts, which possibly increase in relation to increased courtship activity.

**Table 1:** Mean MSP1 and MSP3 amounts in ng/individual (± 1SE, where applicable) for headspace samples and solvent extracts for each sex ratio and mating number.

| Table 1: Mean MSP amounts in ng/individual (± 1SE, where applicable) for each sex ratio and mating number | | | | | |
| --- | --- | --- | --- | --- | --- |
|  |  | Headspace sampling |  | Solvent extracts |  |
| Sex ratio | Mating number | MSP1 | MSP3 | MSP1 | MSP3 |
| 1:3 | 0 | - | - | - | - |
| | 1 | 2198 ± 620 | 920 ± 333 | 2766 ± 495 | 10205 ± 1120 |
| | 2 | 4195 ± 820 | 2490 ± 1120 | 3580 ± 1130 | 12486 ± 3880 |
|  | 3 | 13472 | 8276 | 4817 | 18086 |
| 1:1 | 0 | 1279 | 31 | 3745 | 13880 |
| | 1 | 2280 ± 417 | 897 ± 148 | 3568 ± 478 | 12710 ± 1677 |
|  | 2 | - | - | - | - |
|  | 3 | - | - | - | - |
| 3:1 | 0 | - | - | - | - |
| | 1 | 2495 ± 240 | 850 ± 91 | 2833 ± 842 | 11214 ± 2371 |
| | 2 | 2705 ± 263 | 808 ± 131 | 3179 ± 403 | 11030 ± 1614 |
| | 3 | 2835 ± 89 | 957 ± 151 | 2836 ± 306 | 10999 ± 3460 |

**Table 2:** Summary of models testing for the effects of sex ratio and mating number on MSP1 and MSP3, where male and female age were used as a random factors. Non-significant terms were removed from the full model; hence only significant factors and interactions are listed.

|  |  | Headspace |  |  | Solvent extract |  |  |
| --- | --- | --- | --- | --- | --- | --- | --- |
| Variables | Model terms | Model estimate +/- 1SE | LRT | p | Model estimate +/- 1SE | LRT | p |
| MSP1 | Intercept | -1180.3 +/- 1454.0 |  |  | 2755.0 +/- 988.0 |  |  |
|  | Sex ratio | 1496.9 +/- 638.0 | 0.3 | 0.558 |  |  |  |
|  | Mating number | 3509.8 +/- 886.4 | 5 | 0.03 |  |  |  |
|  | Sex ratio*Mating number | -1205.9 +/- 394.6 | 9.2 | 0.002 |  |  |  |
| MSP3 | Intercept | -1536.0 +/- 932.5 |  |  | 9898.4 +/- 3515.7 |  |  |
|  | Sex ratio | 899.1 +/- 377.8 | 1 | 0.218 |  |  |  |
|  | Mating number | 2473.2 +/- 544.7 | 1 | 0.02 |  |  |  |
|  | Sex ratio*Mating number | -833.8 +/- 273.3 | 11.4 | < 0.001 |  |  |  |

### Do males actively control MSP emission to courtship activity?

We aimed to assess whether males can actively control MSP emission or whether MSP components are emitted passively. We hypothesized that if MSP emission is actively controlled by males, MSP headspace amounts should correlate with courtship activity, though not with general mobility (i.e. walking, flying), because MSP should be emitted specifically when MSP are useful, i.e. when courting females. We contrasted two types of male behaviors that were recorded during mating experiments: male sexual activity as represented by male courtship (fluttering and thrusting) and male general movements (walking, flying) as an internal control. Both MSP1 and MSP3 headspace amounts increased significantly with male courtship activity, while MSP1 and MSP3 headspace amounts decreased significantly when male general movements increased (Figure 4; Table 3). Males thus appear to actively emit MSPs during courtship activity, while simple wing movements produced during flight have an opposite effect on MSP emission. No MSP2 was found to be emitted.

**Table 3:** Summary of models testing for the effect of courtship behaviors and general movements on MSP1 and MSP3 headspace amounts, where male and female age were used as random factors. Non-significant terms were removed from the full model; hence only significant factors and interactions are listed.

| Variables | Model terms | Model estimate +/- 1SE | LRT | p |
| --- | --- | --- | --- | --- |
| MSP1 | Intercept | 1714.6 +/- 1338.3 |  |  |
|  | Courtship behaviors | 2451.7 +/- 916.9 | 7.2 | 0.0007 |
|  | General movements | -1670.9 +/- 655.0 | 6 | 0.014 |
| MSP3 | Intercept | -1186.4 +/- 996.7 |  |  |
|  | Courtship behaviors | 1215.2 +/- 567.9 | 5,8 | 0.016 |
|  | General movements | -1104.4 +/- 421.9 | 7,5 | 0.006 |

LRT = log-likelihood ratio test

**Figure 4:**
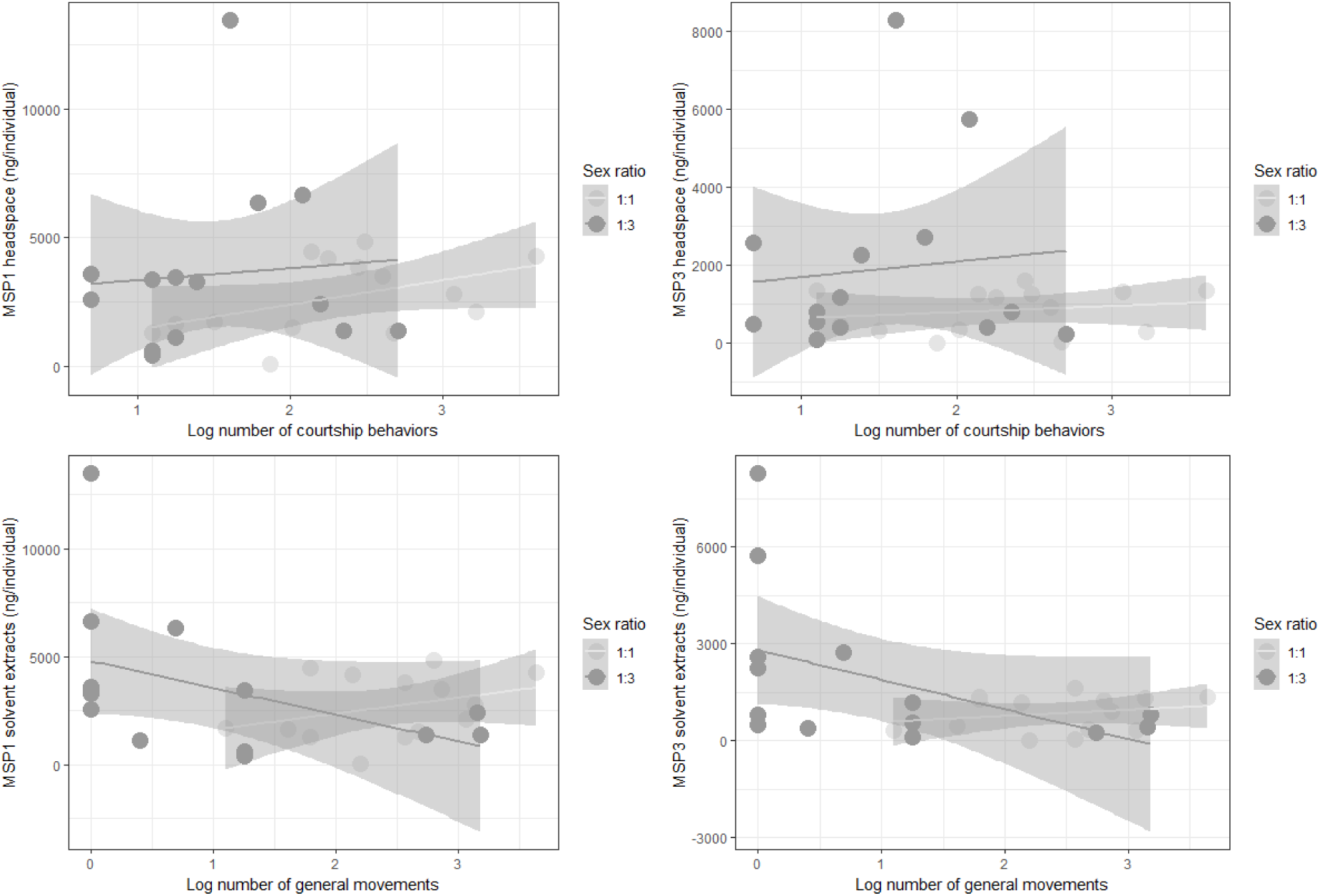
Headspace amounts (in ng/individual) and confidence intervals for each sex ratio in response to increasing courtship activity (top) or general movement (bottom) for MSP1 (left) and MSP3 (right). N = 28 (i.e. treatments where only 1 male was present).

## Discussion

Our results show that solvent extraction, one of the most common methods used to quantify olfactory signals, does not reflect MSP quantities as they are available in the air for female perception. MSP2 and MSP3 indeed displayed strikingly different average amounts when sampled using headspace or solvent extraction, yet MSP1 and MSP3 amounts remained correlated across sampling methods. We also showed that male mating success correlated to increasing amounts of MSP1 and MSP3 when collected using headspace sampling, but not using solvent extracts, and males mated despite the absence of detectable levels of MSP2 in headspace extracts. Absence of a correlation between male solvent extracts and mating success is unlikely due to methodological differences, because male wings were extracted directly after behavioral experiments had ended, as in previous studies on *B. anynana* (Bacquet et al., 2015; Costanzo and Monteiro, 2007; Nieberding et al., 2008, 2012; Prudic et al., 2011). Finally, MSP amounts emitted in the air were found to increase with courtship activity (i.e. wing fluttering), but to decrease in relation to general movements (i.e. walking, flying). Males could thus actively control the emission of MSP when courting females.

How do the discrepancies between headspace and solvent extracts affect our understanding of sexual selection acting on olfactory communication in *B. anynana*? The most striking difference between MSP solvent and headspace extracts was the absence of MSP2 in the latter, whereas on average 1.2 ± 0.05 (1SE) μg/individual was present in solvent extracts. Absence of MSP2 in headspace samples could be due either to technical limitations or to a behavioral decision by males, which were indeed found to control MSP1 and MSP3 emission. Technical limitations are unlikely for several reasons. First, MSP1 (tetradecen-1-ol) was detected in large amounts in headspace samples, and this fatty acid derived component is a long-chain molecule like MSP2 (hexadecanal); hence the polarity of the two molecules, and their adsorption on/desorption from the headspace column, are similar. MSP2 should thus have been found in headspace samples if it had been emitted by males. Second, we used different types of cartridges (Tenax TA, Super Q, Poropak, HayeSep and Silice), as well as two cartridges in series (Tenax-TA) during pilot experiments. Third, a range of flow rates were used in pilots for headspace collection, ranging from 75 mL min^-1^ up to 800 mL min^-1^. We further used males of different ages and densities of up to ten males. None of these pilot experiments led to collection of even trace amounts of MSP2, after control by GC-MS (data not shown). Absence of MSP2 in headspace samples thus suggests that MSP2 emission is actively controlled by males and that the experimental environment used did not elicit active emission of MSP2.

A plausible explanation for the lack of MSP2 production by males may be the limited volume of the entrainment chamber (1.8 L) and long experimental durations (22.5 hours). The role of limited cage size and repeated male-female interactions has recently been shown to artificially strengthen the relative importance of male-male competition over expression of female mate preference for determining mating success in this butterfly (Gauthier et al, 2015; Holveck and Nieberding 2017; Nieberding and Holveck 2018). Females were not able to escape this unnaturally small arena, providing males with an overall high chance of mating (Gauthier et al, 2015; Holveck and Nieberding 2017; Nieberding and Holveck 2018). Such a small-sized and crowded environment may lead males to not invest in the emission of the MSP2 component (Nieberding and Holveck 2018, and refs therein). Absence of MSP2 in headspace extracts could also be explained if MSP2 was an arrestant pheromone, i.e. a signal emitted by flying males to stimulate landing by females before males start their land-based courtship sequence (Clearwater, 1972). Further experiments using a range of cage volumes, including much larger cages allowing flight and escape to take place, coupled to headspace extractions, will be needed to tease these two explanations apart.

The comparison of headspace and solvent extracts also revealed that the relative amounts of MSP1 and MSP3 were inverted between the two methods of quantification: there was about two and a half times more MSP1 than MSP3 in the air, while five times less MSP1 than MSP3 is usually found in solvent extracts (Nieberding et al., 2008, 2012). As MSP1 and MSP3 amounts quantified from headspace, but not solvent, extracts correlated with mating success, our results suggest that headspace sampling of MSP components is a better proxy of what females perceive to assess male quality than solvent extracts, when both methods of quantification are compared with a robust statistical approach. Compared to experiments using headspace extracts, a much larger sample size thus appears needed to spot differences in mating success due to variation in MSP levels when solvent extracts are used (e.g. experiments involving hundreds of males, as in Nieberding et al, 2008, 2012).

We can reasonably conclude that headspace extracts are more reliable estimates of olfactory signals as they are perceived, and under sexual selection by females, compared to solvent extracts. Our results highlight that our understanding of how sexual selection shapes olfactory sexual communication in this model butterfly may be biased by our sampling methodology. Relative proportions of MSP components are known to be of great importance for species recognition in sexual interactions of many Lepidoptera (Groot et al., 2006, 2016). These relative proportions appear to be inversed regarding MSP1 and MSP3 between methods of quantification. In addition, sexually selected traits involved in assessing male quality are usually under strong directional selection for increasing amounts (e.g. Rodríguez et al., 2013). Hence, we may have been biased in previous studies with *B. anynana* by believing that MSP3 was possibly under strongest directional sexual selection as this MSP was present in highest amounts on male wings. While MSP3 amount did increase in some (but not all; Van Bergen et al, 2013) studies with mating success, this may simply be due to the fact that MSP3 amount correlates to MSP1 amount (Nieberding et al., 2012; this study). This study further suggests that the most important MSP component for mating success, MSP2 (Nieberding et al, 2012; Dion et al, 2017), may be under active control for emission by males and the environment in which males are tested may matter for MSP2 emission. We may thus underestimate the role of chemical communication in mating success and sexual selection, and particularly of MSP2 amounts in *B. anynana*, by using unnatural, laboratory-based, environmental conditions (Holveck and Nieberding 2017; Miller & Svensson, 2014; Nieberding & Holveck, 2018). Although pheromones that are present in the air may not equate to what is perceived by females (e.g. because pheromone perception may depend on the number of olfactory receptors dedicated to each MSP component on the antenna), this study reveals potential limitations of using solvent extracts of olfactory tissues and organs as proxies for olfactory communication as it evolves under sexual selection.

## Author contributions

BV analysed the data, wrote and edited the manuscript; IAND, SH, GL and CMN developed the experimental methods; IAND, SH, FL and PMBB collected data; IAND produced technical illustrations; GL discussed and edited the manuscript; CMN conceived and designed the research, analysed the data, wrote and edited the manuscript.

## Acknowledgments

We would like to thank two anonymous reviewers for their helpful suggestions on an earlier draft of this paper. We are further grateful to Christophe Pels for assistance with the butterfly rearing. This is BRC publication 415 of the Biodiversity Research Centre. This paper was posted on bioRxiv prior to publication (Visser et al., 2018).

## Data availability

Data is available as supplementary information.

---

**Supplementary figure 1:**
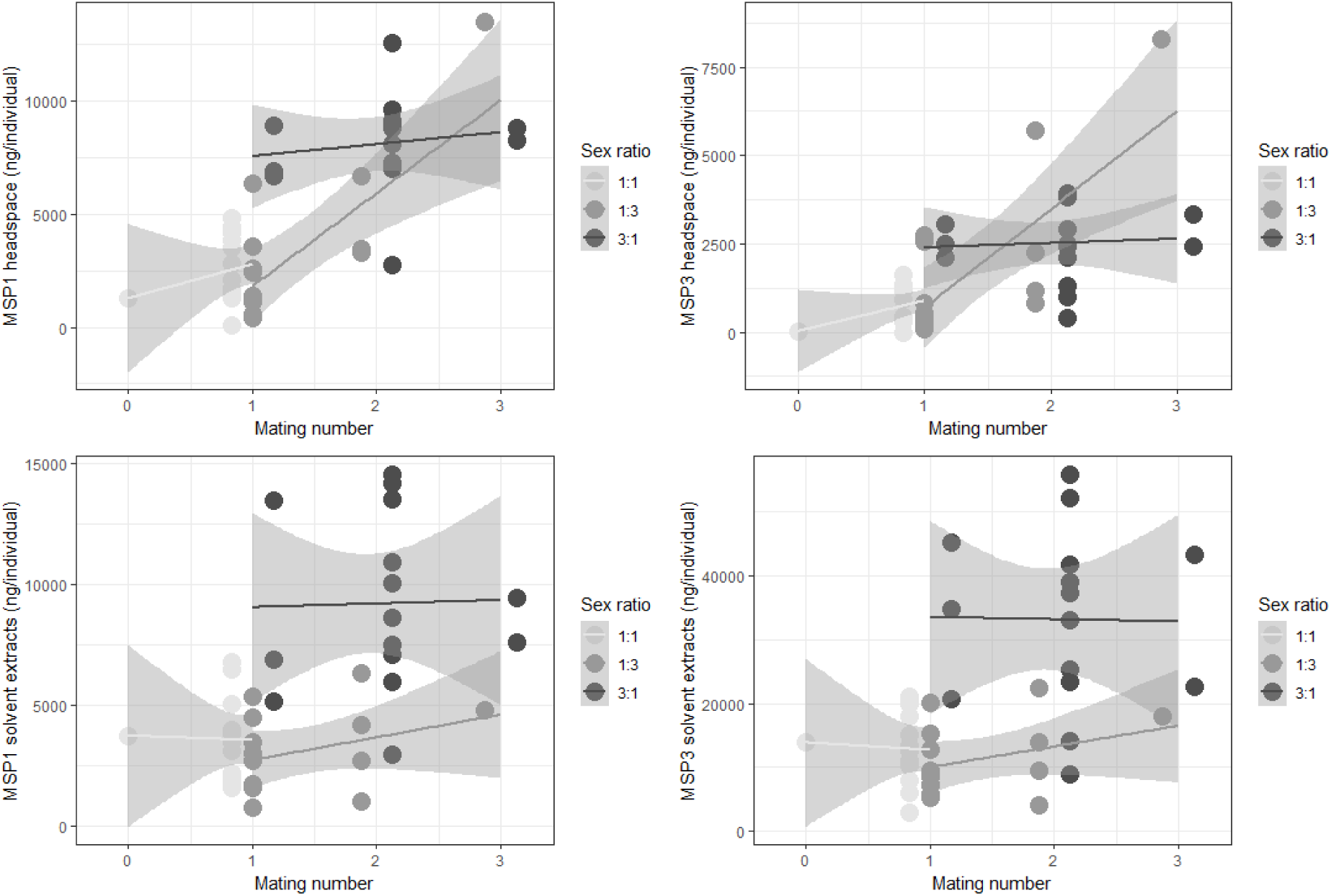
MSP1 (left) and MSP3 (right) headspace amounts (in ng/individual; top) and solvent extract amounts (in ng/individual; bottom) for each mating number (X-axis) and sex ratio treatment.

